# YbeY is required for ribosome small subunit assembly and tRNA processing in human mitochondria

**DOI:** 10.1101/2019.12.12.874305

**Authors:** Aaron R. D’Souza, Lindsey Van Haute, Christopher A. Powell, Pedro Rebelo-Guiomar, Joanna Rorbach, Michal Minczuk

## Abstract

Mitochondria contain their own translation apparatus which enables them to produce the polypeptides encoded in their genome. The mitochondrially-encoded RNA components of the mitochondrial ribosome require various post-transcriptional processing steps. Additional protein factors are required to facilitate the biogenesis of the functional mitoribosome. We have characterised a mitochondrially-localized protein, YbeY, which interacts with the assembling mitoribosome through the small subunit. Loss of YbeY leads to a severe reduction in mitochondrial translation and a loss of cell viability, caused by less accurate mitochondrial mt-tRNA^Ser(AGY)^ processing from the primary transcript and an accumulation of immature mitochondrial small subunit. Our results suggest that YbeY performs a dual function in mitochondria coupling tRNA processing to mitoribosome biogenesis.

**Issue Section:** Nucleic Acid Enzymes

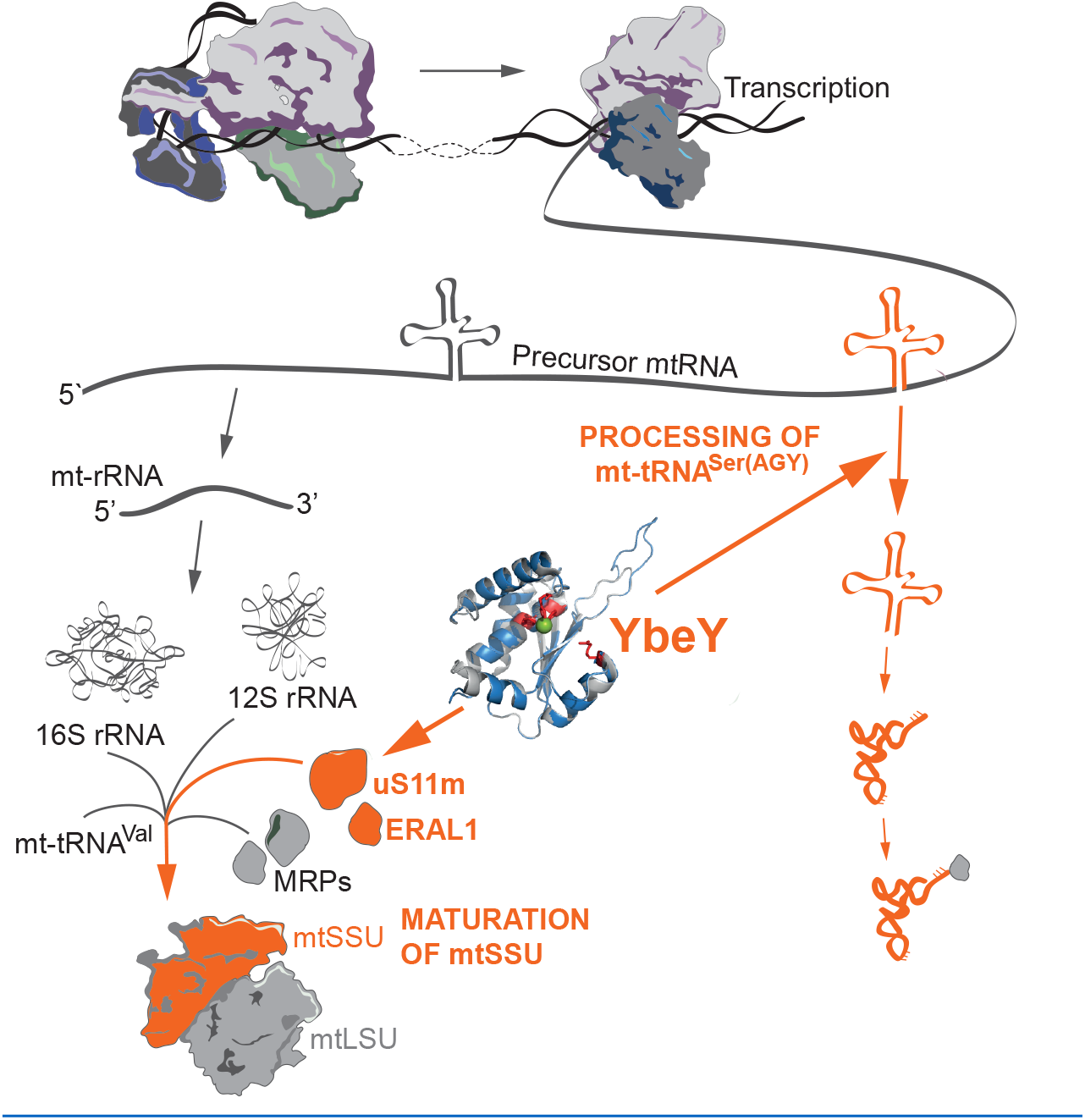

## INTRODUCTION

Oxidative phosphorylation (OxPhos) is one of the major functions of mitochondria. A majority of the components of the OxPhos complexes are encoded in the cell nucleus and imported into mitochondria upon translation in the cytoplasm. However, thirteen structural polypeptides of the OxPhos complexes are translated within mitochondria. These polypeptides are encoded in the human mitochondrial genome (mtDNA) along with the twenty-two tRNAs and two rRNAs that make up the RNA components of the intra-organellar translation apparatus.

The mtDNA is transcribed bidirectionally, generating long polycistronic RNA strands. In the majority of cases, mRNAs and rRNAs in the precursors are flanked by tRNA molecules (1,2). The excision of these mt-tRNAs is performed by endonucleolytic enzymes RNase P at the 5’ end, and RNase Z (ELAC2) at the 3’ end (3,4). This releases the mt-mRNAs and mt-rRNAs, with the latter undergoing further maturation and assembly into the mitochondrial ribosome.

The mammalian mitoribosome consists of a large (mt-LSU) and small subunit (mt-SSU), and is composed of approximately 80 nuclear-encoded proteins, two mt-rRNAs and one mt-tRNA (5,6). The assembly of the mitoribosome involves the chemical modification of the mt-rRNAs, spontaneous RNA-protein and RNA-RNA interactions and RNA folding, and requires the assistance of various assembly factors. These factors include, but are not limited to, GTPases and ATP-dependent RNA helicases. The hydrolysis of NTPs by these enzymes ensure that the mt-rRNA does not fall into conformational traps. For example, GTPBP10 (also called ObgH2) (7,8), OBGH1 (also called GTPBP5) (9) and MTG1 (also called GTPBP7) (10) are three conserved GTPases that assist in the assembly of mt-LSU, and C4orf14 (11–13) and ERAL1 (14,15) are two conserved GTPases that are involved with the biogenesis of mt-SSU. RNA helicases such as DDX28 (16) and DHX30 (17) unwind mt-rRNA coupled to the hydrolysis of ATP. Additional factors such as MALSU1 form a trimeric complex with L0R8F8 and mt-ACP at the intersubunit interphase preventing the association of the immature subunits during monosome formation (18,19). The early stages of mitoribosomal assembly occur co-transcriptionally in the nucleoid (20). The assembly of mt-LSU can initiate on the 12S-16S mt-rRNA precursor molecule, but not the mt-SSU (21). The initiation and progression of small subunit assembly may be inhibited until a certain stage of early mt-LSU assembly is complete (8,22). Then, subsequent steps of mitoribosome biogenesis are continued in distinct foci called RNA granules (23,24).

In bacteria, the ribosomal RNAs are co-transcribed as a single operon. The pre-rRNA undergoes the removal of transcribed spacer regions before the rRNA is ready for assembly. The bacterial homolog of human YbeY is a single strand-specific endoribonuclease that is responsible for the 3’ end processing of the SSU rRNA (25–28). Depletion of the protein leads to accumulation of precursors, the loss of the SSU and a subsequent translation defect (29,30). In addition, the protein also plays a role in LSU rRNA and 5S rRNA maturation (26). However, recent work suggests that, unlike other nucleases that can use immature small subunits containing precursor 16S rRNA as substrates for nucleolytic activity, the endonucleolytic activity of purified bacterial YbeY has not been demonstrated *in vitro*. Even the addition of various combinations of known YbeY interactors, ribosome components and initiator tRNA did not lead to expected YbeY-specific RNA cleavage events (27,31). Furthermore, it has also been implicated in the degradation of defective and misprocessed SSU rRNA and, as such, plays a role in ribosome quality control, in conjunction with RNase R (26,31,32).

Very little is known about the human orthologue of bacterial YbeY. Unlike bacteria, the 16S and 12S mt-rRNA components are flanked by mt-tRNAs, and their excision by RNase P and ELAC2, without the need for further end processing, is sufficient for assembly into the mitochondrial ribosome (33,34). Therefore the function of mammalian YbeY is expected to be substantially different. Recent work demonstrated that the introduction of human YbeY into bacteria lacking the homolog caused partial rescue of phenotype (35). However, the exact function of YbeY in mammalian mitochondria has so far not been investigated.

We and others have recently shown that defects in the structural components of mitoribosome and its assembly factors can lead to a human disorder of mitochondrial respiration (36). The RNase activity of YbeY from various bacterial species (26,28,37), plant chloroplasts (38) and the mammalian mitochondria have been previously demonstrated. Therefore, we have focussed on the functional significance of the protein in the mitochondria (35). Here, we show that YbeY is a mitochondrially-localised protein. We characterise YbeY-deficient HEK293T cell lines and YbeY knockout Hap1 cell lines. We demonstrate that the loss of the protein leads to a severe deficiency in mitochondrial translation. We also show that YbeY has a dual function: (i) YbeY plays a role in tRNA^Ser(AGY)^ processing at both the 5’ and 3’ ends and (ii) interacts with the mt-SSU component uS11m. The loss of YbeY causes a decrease in the abundance of mt-SSU components and an increase in mt-LSU component abundance. Furthermore, in YbeY knockout cells, mt-SSU and mt-LSU fail to effectively assembly into a monosome. These findings on the role of YbeY provide new insights into the mitoribosome assembly process and, in the long-term, can help pave the way for the development of future mechanism-based therapies for mitochondrial diseases.

## RESULTS

### YbeY is a mitochondrial protein with a conserved endonuclease domain

Human YbeY is an uncharacterised protein that is encoded by the *C21orf57* gene. It belongs to the UPF0054 protein family (PFAM accession number: PF02130) which contains a highly conserved zinc ion-coordinating consensus motif H3XH5XH (39). The bacterial homolog is an active endoribonuclease and contains the RNA-binding Arg59 residue and the histidine triad (26,32). Sequence alignment from multiple species (**Figure 1** and **Supplementary Figure S1**) and homology modelling of human YbeY, using the *E. coli* protein as the template, revealed strong conservation, especially of the aforementioned active site residues, suggesting that human YbeY is an active RNase (**Figure 1B**). In addition, the previous analysis showed that *E. coli* Asp85 (Asp90 *H. sapiens*), found in the beta-sheet outside the active site and required for the interaction of YbeY with the ribosomal small subunit component, S11, is also conserved. Mutation of these residues has been shown to disrupt nuclease activity of the protein and rRNA maturation (35) (**Supplementary Figure 1**). Further comparison of the two homologs also shows that human YbeY has a nine amino acid insertion (Gly75-Pro83) between two conserved beta sheets which is conserved in other eukaryotic species (**Supplementary Figure 1**) and may be responsible for additional protein interactions and additional functions in the mitochondria (**Figure 1B**) (35).

**Figure 1.**
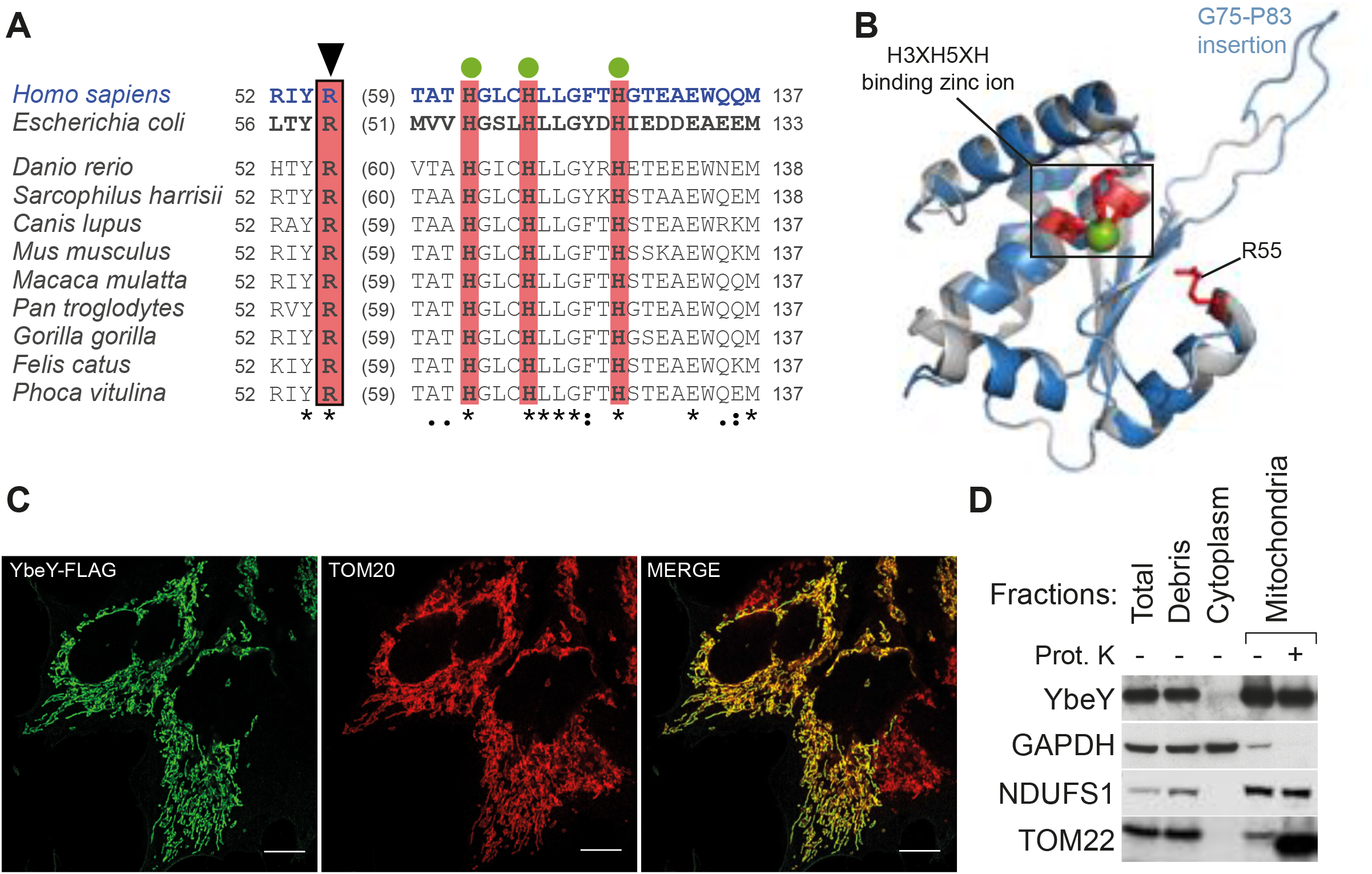
Sequence/structure conservation and mitochondrial localisation of human YbeY. (A) Alignment of protein sequences of the orthologues of YbeY. The catalytically-important RNA-binding arginine is indicated with a box and a triangle above. Green circles indicate histidine residues from the zinc ion-binding H3H5XH motif. Colons and dots denote chemical similarity between the sequences, asterisks indicate identical residues. (B) Homology modelling of the human YbeY using *E. coli* YbeY as a template (PDB: 1XM5) superimposed on each other: human YbeY (blue), bacterial YbeY (grey), conserved catalytically important residues (red) and zinc ion (green). Eukaryote-specific insertion (Gly75-Pro83 in *H. sapiens*) is indicated. (C) Flag-tagged YbeY was transiently expressed in HeLa cells and visualised using anti-flag antibody (green) and the mitochondrial outer membrane translocase, TOM20 (red). Co-localisation is shown in yellow. The white scale bar at the bottom right depicts 10 μm. (D) Subcellular fractionation of HEK293T cells shows YbeY protein in the same fraction as NDUFS1 (mitochondrial inner membrane) and TOM22 (mitochondrial outer membrane, truncated by proteinase K), but not with GADPH (cytoplasm)

YbeY has been previously predicted to be mitochondrially localised by large-scale databases, MitoCarta 2.0 and MitoMiner 4.0 (40,41). Furthermore. large-scale CRISPR/Cas9 screen has revealed that YbeY is vital for OXPHOS function (42). In accordance with these results, *in silico* analysis of the subcellular localisation of the protein by multiple online tools predicted a high probability of mitochondrial localisation: Predotar (43) – 0.87, TargetP (44) – 0.86, and Mitoprot (45) – 0.96. To confirm these predictions, we used immunostaining that showed a co-localisation of transiently-expressed flag/strep2-tagged YbeY to the mitochondria in HeLa cells (**Figure 1C**). We also used subcellular fractionation and immunoblotting to localise endogenous YbeY in the cell. YbeY co-fractionated with inner membrane-localised Complex I subunit, NDUFS1 (**Figure 1D**). To assess, if YbeY co-localises within mitochondrial RNA granules (MRGs), where processing of nascent RNA occurs, immunocytochemistry was performed using antibodies against a MRG marker GRSF1 (46). While the MRGs appeared as distinct puncta, YbeY was distributed throughout the mitochondria (**Supplementary Figure S2**). Taken together, this confirms that YbeY is a mitochondrial protein but does not co-localise within MRGs.

### YbeY is required for mitochondrial translation

In order to investigate the role of YbeY, we inactivated its expression in human cells. YbeY-deficient HEK293T cell lines were generated using zinc finger nuclease targeting exon 2, whereas YbeY knockout Hap1 cells were created by CRISPR/Cas9. The hemizygous HEK293T knockout – hereby referred to as YbeY (−/m) – contained three out-of-frame deletions and one allele with a 12 bp deletion and a change in 4 amino acids (**Supplementary Figure S3A**). In the heterozygous HEK293T mutant (YbeY (+/−)), three alleles contained indels which led to premature stop codons, while one allele contained an in-frame deletion leading to a loss of two amino acids (**Figure 3B**). We also used a near haploid, Hap1, YbeY knockout (YbeY (−/−)) cell line that contained a 13 bp insertion (**Supplementary Figure S3C**) which produced a premature stop codon and leads to an 80 amino acid truncated protein. All three YbeY-deficient cell lines exhibited severely depleted levels of endogenous protein (**Figure 2A, B**).

**Figure 2.**
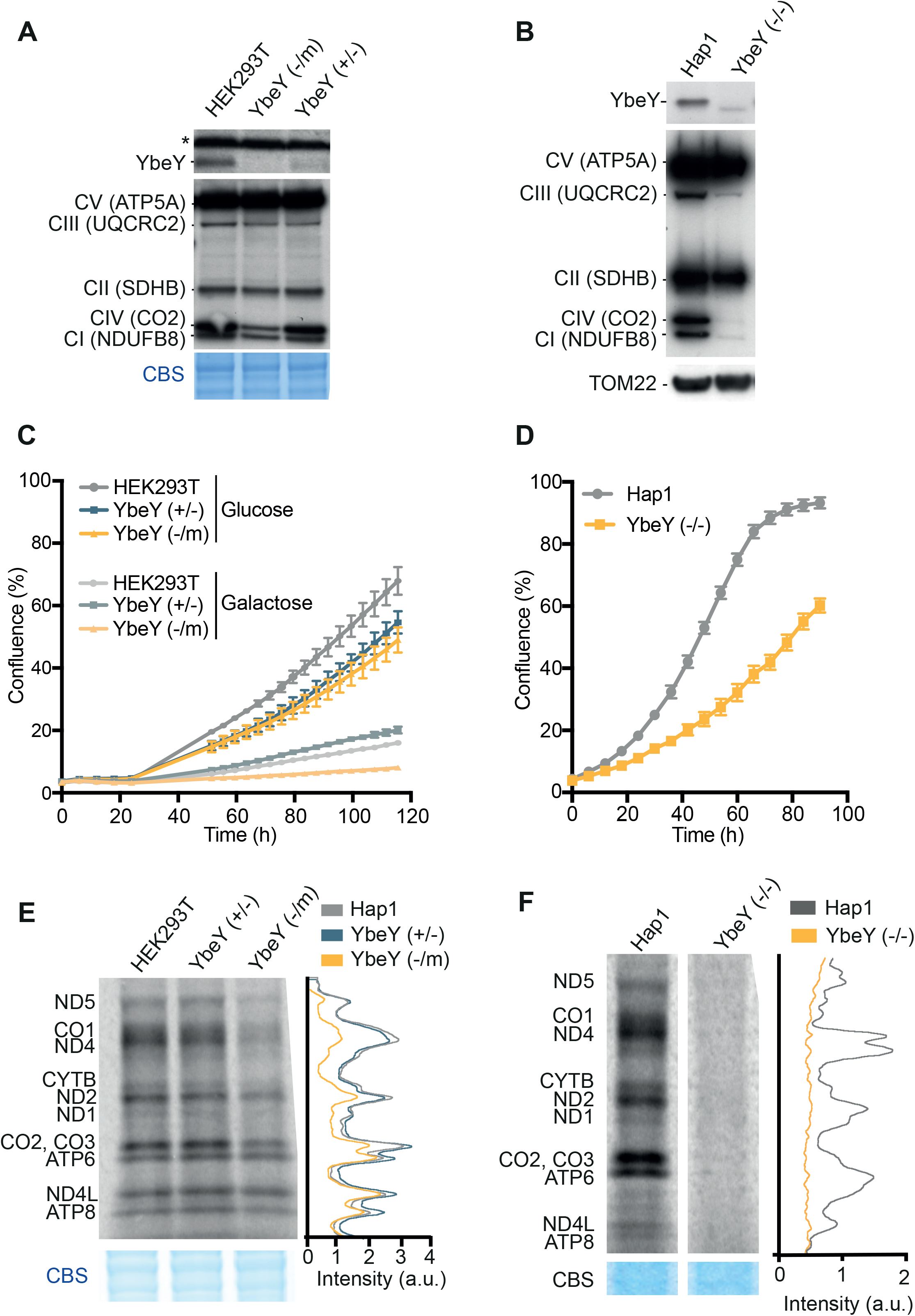
Reduced mitochondrial gene expression and growth in cells lacking YbeY. (A) Analysis of steady state levels of endogenous YbeY and OXPHOS subunits in HEK293T parental, cell that 1 functional copy of *YBEY* gene (+/−) and YbeY-deficient (−/m) cells. Coomassie Blue staining (CBS) was used as loading control. (B) Analysis of steady state levels of endogenous YbeY and OXPHOS subunits in Hap1 parental and YbeY-deficient (−/−) cells. TOM22 was used as a loading control. (C) Growth of HEK293T parental and YbeY-deficient cells in glucose and galactose media. Representative experiment is shown where cells were plated in triplicate (mean ±1 SD). (D) Growth of Hap1 parental and YbeY knockout cells in high glucose IMDM media containing 20% FBS. Representative experiment is shown where cells were plated in triplicate (mean ±1 SD) (E) Metabolic labelling of mitochondrial translation products with [^35^S]-methionine in HEK293T parental cells, YbeY (+/−) and YbeY (−/m) cell lines. Coomassie Blue staining (CBS) was used as loading control. (F) *De novo* protein synthesis of Hap1 parental cells and YbeY (−/−) cells. Coomassie Blue staining (CBS) was used as loading control.

**Figure 3.**
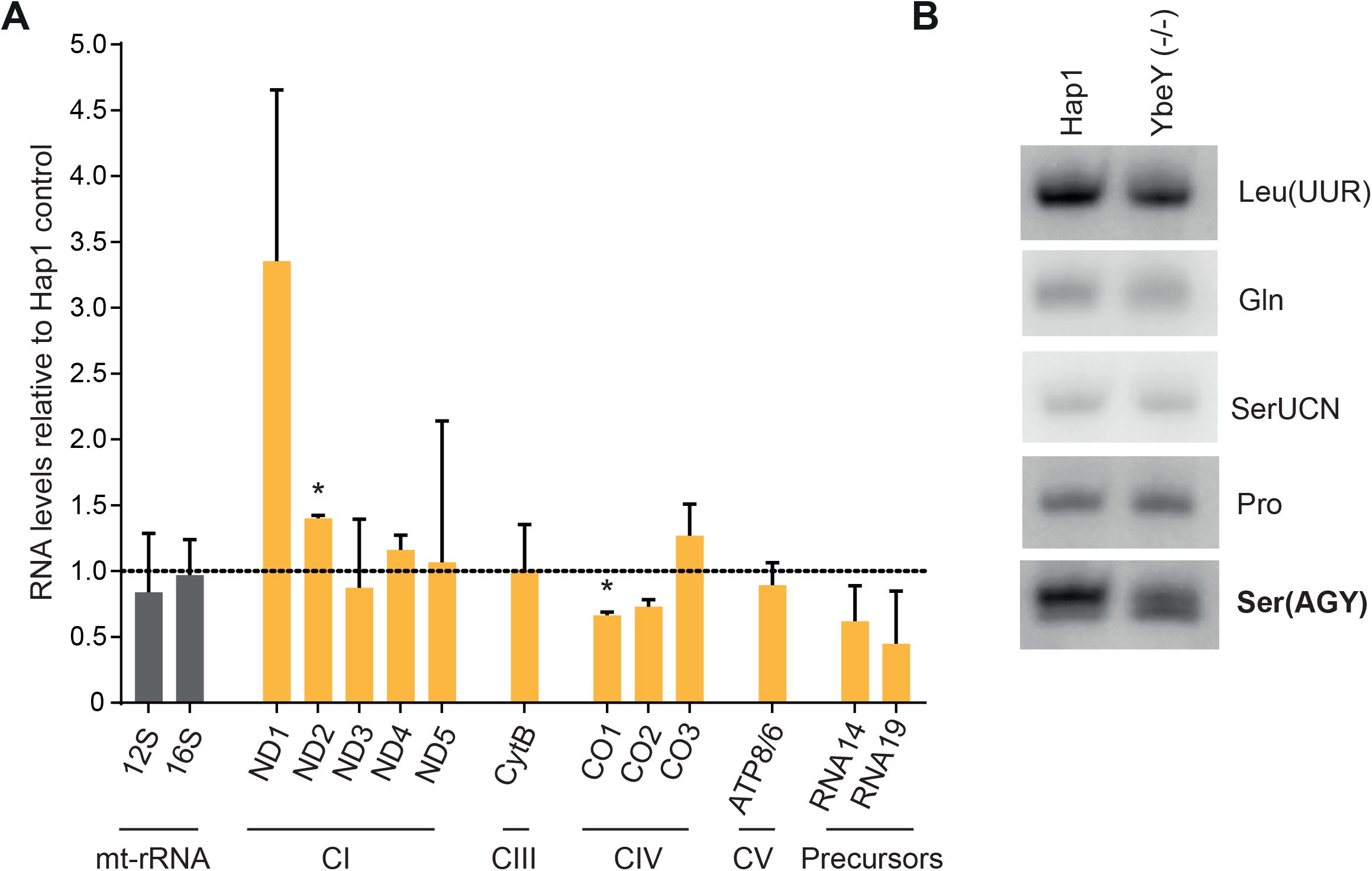
Steady state levels of mitochondrial RNA in YbeY depleted cells. (A) Steady-state levels of mitochondrial RNA quantified by northern blotting in Hap1 control and knockout cells. 5S RNA was used as a loading control. Quantification of steady state levels was performed using Image J and normalised to values from Hap1 control cells. *P<0.05, One sample t-test, Error bars = 1 SD (B) High resolution northern blot analysis of mitochondrial tRNAs from Hap1 control and YbeY knockout cells.

Next, we intended to study the OXPHOS function and mitochondrial translation upon ablation of YbeY. Western blot analysis showed a strong reduction in the steady state levels of Complex I and IV subunits in YbeY(−/m) cells (**Figure 2A**). A much stronger effect was observed in the complete knockout cell line where, in addition to Complex I and IV, a Complex III subunit was also depleted, suggesting a defect in synthesis of the OXPHOS subunits in the absence of YbeY. Consistent with this result, cell growth was compromised when YbeY-deficient HEK293 cells were grown in media containing galactose as the sole carbon source, forcing them to rely on mitochondrial ATP production (**Figure 2C**). The growth of Hap1 YbeY-deficient cells was compromised on standard, glucose-containing medium while the cells failed to grow on medium containing galactose as the sole carbon source (**Figure 2D**). The effect of YbeY depletion on mitochondrial translation was assessed using *in vivo* incorporation of [^35^S]-methionine into newly synthesised proteins. A global mitochondrial translation defect was observed in the YbeY(−/−) cells, while a reduction of protein synthesis was observed in the YbeY(−/m) cells (**Figure 2E, F**). Moreover, in YbeY-deficient HEK293T cells, the higher molecular weight translation products were affected to a greater degree than the lower molecular weight products (**Supplementary Figure S4A**). This loss of translation was mitochondria-specific and not caused as a by-product of reduced cytoplasmic translation (**Supplementary Figure S4B**). Taken together, these result show that YbeY is indispensable for mitochondrial translation.

### YbeY is required for accurate mt-tRNA^Ser(AGY)^ processing

In *E. coli*, YbeY is required for endonucleolytic processing of the rRNA precursors (32). Thus, we set out to investigate if the endonucleolytic function of human YbeY has a role in controlling mitochondrial transcript steady state level. Northern blot analysis showed that there was no difference in the levels of the 12S and 16S mt-rRNAs (**Figure 3A, Supplementary Figure S5A)**. Any observed increase in mt-mRNAs coding for Complex I subunit ND1 and ND2 steady state levels and a decrease in mt-mRNAs encoding Complex IV subunit CO1 and CO2 (**Figure 3A, Supplementary Figure S5A**) was not attributed to YbeY deficiency, as a similar compensatory phenotype has been often observed in cells responding to defective mitochondrial translation (47–50).

Further analysis of mitochondrial tRNA using high resolution northern blotting showed a potential steady-state decrease and/or processing defect of mt-tRNA^Ser(AGY)^ relative to other mt-tRNAs analysed (**Figure 3B**). Given that bacterial YbeY is predominantly involved in rRNA end processing, we wanted to investigate the effects of YbeY knockout on the 5’ and 3’ ends of the mt-rRNAs and mt-tRNAs to see if YbeY plays a similar role in the mammalian mitochondria. To do so, we used circularisation RT-PCR to ligate the 5’ end of the RNA molecule to the 3’ end and sequence the junction. No significant difference was observed in the 5’ and 3’ processing of mt-tRNA^Ile^, mt-tRNA^Leu(UUR)^, 12S mt-rRNA and 16S mt-rRNA when Hap1 parental cells and YbeY knockout cells were compared (**Figure 4A, B**). However, in YbeY knockout Hap1 cells, only ~1 % of the mt-tRNA^Ser(AGY)^ molecules were correctly cleaved from the primary transcript and modified to add the 3’ CCA, as compared to the ~85 % in the control Hap1 cells (**Figure 4A**). Sequence analysis of the mt-tRNA processing in the YbeY knockout cells revealed that only ~ 20 % of mt-tRNA^Ser(AGY)^ were correctly processed from the 5’ terminus and ~64 % from the 3’ end (**Figure 4C, D**). This confirms that YbeY is indispensable for the correct end processing of mt-tRNA^Ser(AGY)^.

**Figure 4.**
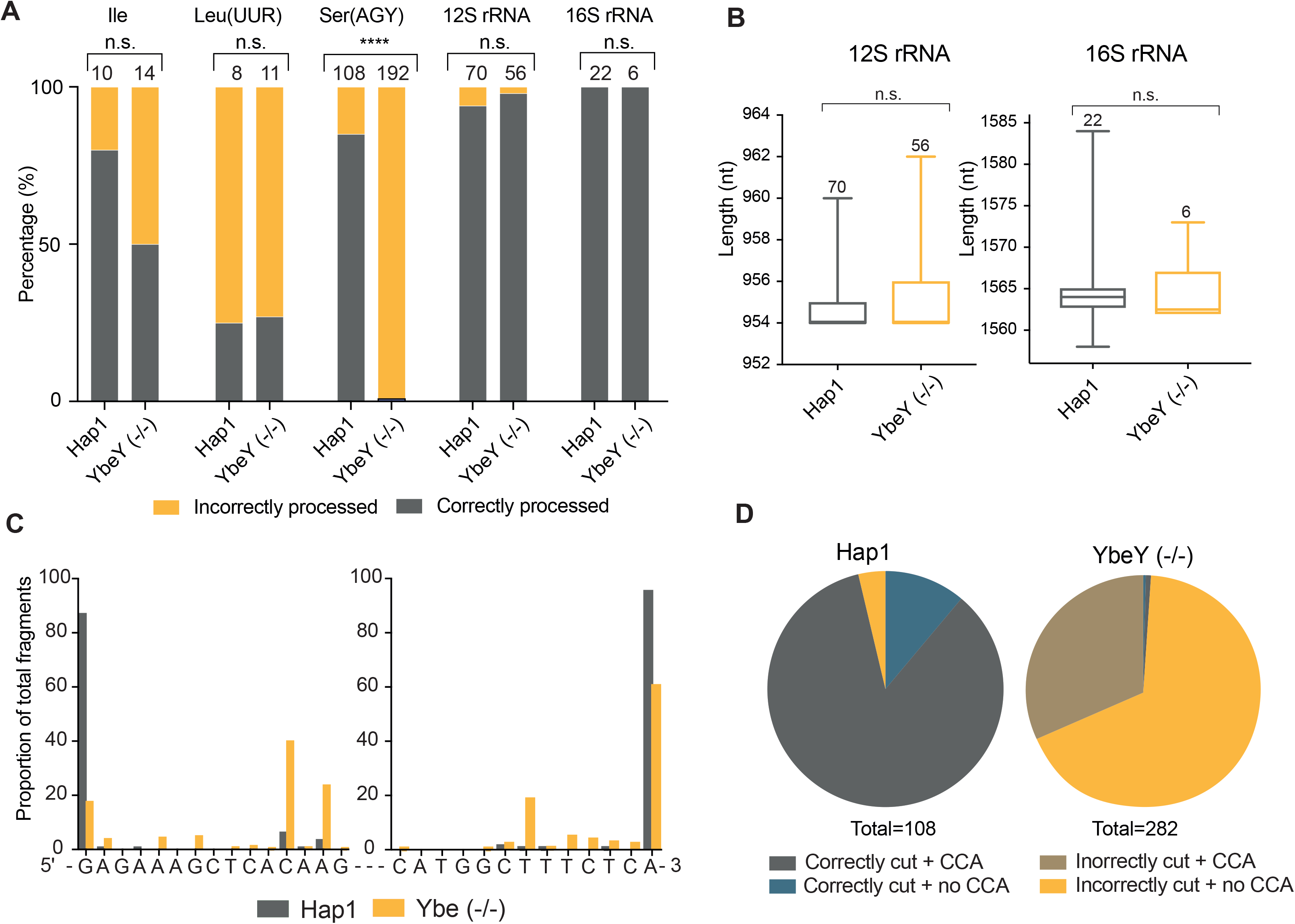
YbeY depletion causes defects in mt-tRNA^Ser(AGY)^ end processing. (A) Results of circularisation RT-PCR analysis of mt-tRNA^Ile^, mt-tRNA^Leu(UUR)^, mt-tRNA^Ser(AGY)^, 12S mt-rRNA and 16S mt-rRNA from Hap1 control and YbeY knockout cells. DNA bands obtained in the circularisation RT-PCR reaction were analysed by Sanger sequencing upon cloning. Number of clones analysed in each sample is given above each bar. Analysis of significance was calculated using Fisher’s exact test. ****p<0.0001 (B) The length of the 12S and 16S mt-rRNAs calculated from the 5’ and 3’ ends. The lines indicate the minimum and maximum values and the edges of the boxes indicate the upper and lower quartile. The middle line indicates the median. The number of data points are written above the box and whisker plots. Significance was assessed using an unpaired Student’s T-test (n.s., not significant). (C) Mapping of the sequence of the 5’ and 3’ ends of circularised mt-tRNA^Ser(AGY)^ in Hap1 parental and YbeY knockout cells. The 5’ and 3’ ends were plotted independently as a percentage of the total 5’ and 3’ reads sequence, respectively. (D) Proportion of mt-tRNA^Ser(AGY)^ that are correctly or incorrectly cleaved from the primary transcript at the expected ends and modified or not to have a 3’ CCA.

### YbeY interacts with mitochondrial ribosomal components

In order to study a potential interaction of human YbeY with the mitoribosomal subunits, we used western blot analysis after sucrose gradient fractionation. A majority of YbeY was found in the “free” fractions (fractions 1-4). However, a small proportion co-migrated with the mt-SSU, suggesting potential weak interaction of YbeY with this subunit (**Figure 5A).** Next, in order to identify interactors of human YbeY with a higher sensitivity, we used quantitative SILAC-based immunoaffinity purification of flag-tagged YbeY. This experiment identified various mt-SSU and mt-LSU components being pulled-down with YbeY. The strongest interactor of YbeY appeared to be uS11m. Other interactors of YbeY included ERAL1 and mitoribosomal proteins such as uL2m, uL4m, bL20m, bL21m, mL42, mL43, mL50, mL64 and mS26 (**Figure 5B, Supplementary Figure S6**). The interaction with uS11 and ERAL1 is highly conserved in bacteria and vital for the maturation of the small subunit (51). These results show conserved interactions of YbeY with the ribosome and, hence, a potentially conserved role in ribosome biogenesis between bacteria and human mitochondria.

**Figure 5.**
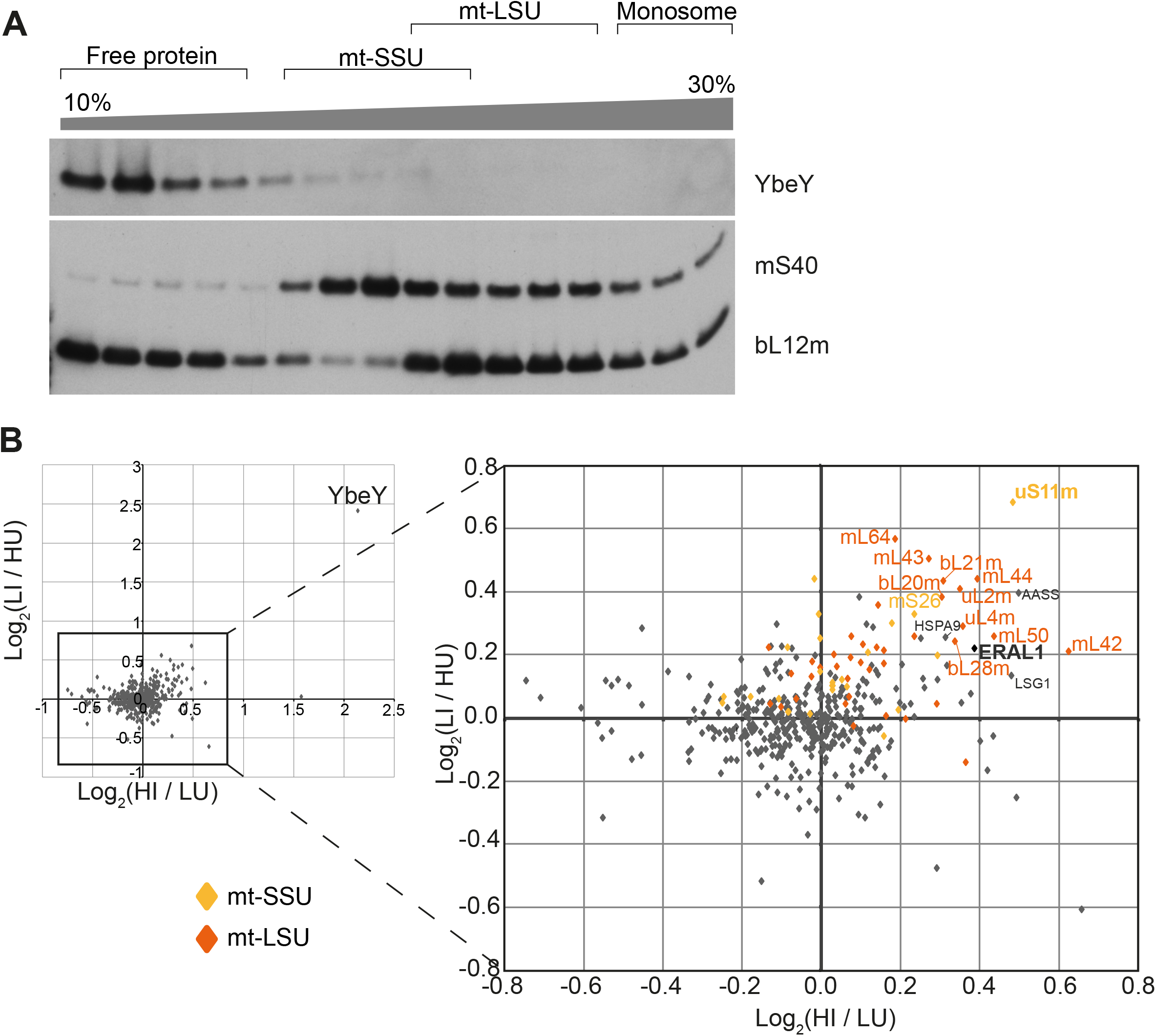
Human YbeY interacts with uS11m. (A) Analysis of the interaction of endogenous YbeY with the mitochondrial ribosome using isokinetic sucrose gradient fractionation and western blotting. Fractions containing unassembled free protein, the subunits at various stages of assembly, and the associated mitoribosome are indicated above. Antibodies against mS40 and bL12m were used to identify the fractions containing the small and large subunits respectively. (B) Immunoaffinity purification of flag-tagged YbeY. The log_2_(ratio of peptide abundance in induced cells to peptide abundance in uninduced cells) were plotted on the X and Y axes from reciprocally labelled experiments. Proteins in the top right quadrant are those that were enriched in the YbeY-overexpressing cell line and proteins in the lower left quadrant are those that are more abundant in the control cell line. Mt-SSU components (yellow diamonds), mt-LSU components (orange diamonds) and other proteins (grey diamonds).

### Loss of YbeY impairs mitoribosome assembly

In *E. coli*, YbeY is responsible for ribosomal quality control. With the assistance of exonuclease RNase R, bacterial YbeY degrades rRNA from late stage ribosome assembly intermediates with defective small subunits (26,32). Hence, we intended to investigate the effect of YbeY depletion on the mitoribosome integrity. Western blot showed a decrease in the steady state levels of mt-SSU components, while the steady-state levels of mt-LSU remained largely unchanged, as compared to a control (**Supplementary Figure S7A**). To further verify this, we performed western blot analysis after sucrose gradient fractionation. Again, we observed a decrease in abundance of the assembled mt-SSU, with a large proportion of mt-SSU (measured by the abundance of uS17m) being present in the earlier ‘free protein’ fractions (**Supplementary Figure S7B-C**). To quantitatively assess changes in composition and integrity of the mitoribosome in the YbeY knockout cells, we used isokinetic sucrose gradient fractionation in combination with SILAC and quantitative mass spectrometry. These experiments confirmed that mt-SSU was strongly decreased in the YbeY knockout cells (**Figure 6A, C**), while mt-LSU was more abundant in the knockout cells relative to the control (**Figure 6B, D**). A comparison of the ratio of assembled mt-SSU and mt-LSU subunits to the monosome implies that in cells lacking YbeY, a larger proportion of both subunits is not assembled into the monosome (**Figure 6E**). Taken together these experiments indicate a problem in mt-SSU assembly/stability preventing correct formation of the monosome in mitochondria.

**Figure 6.**
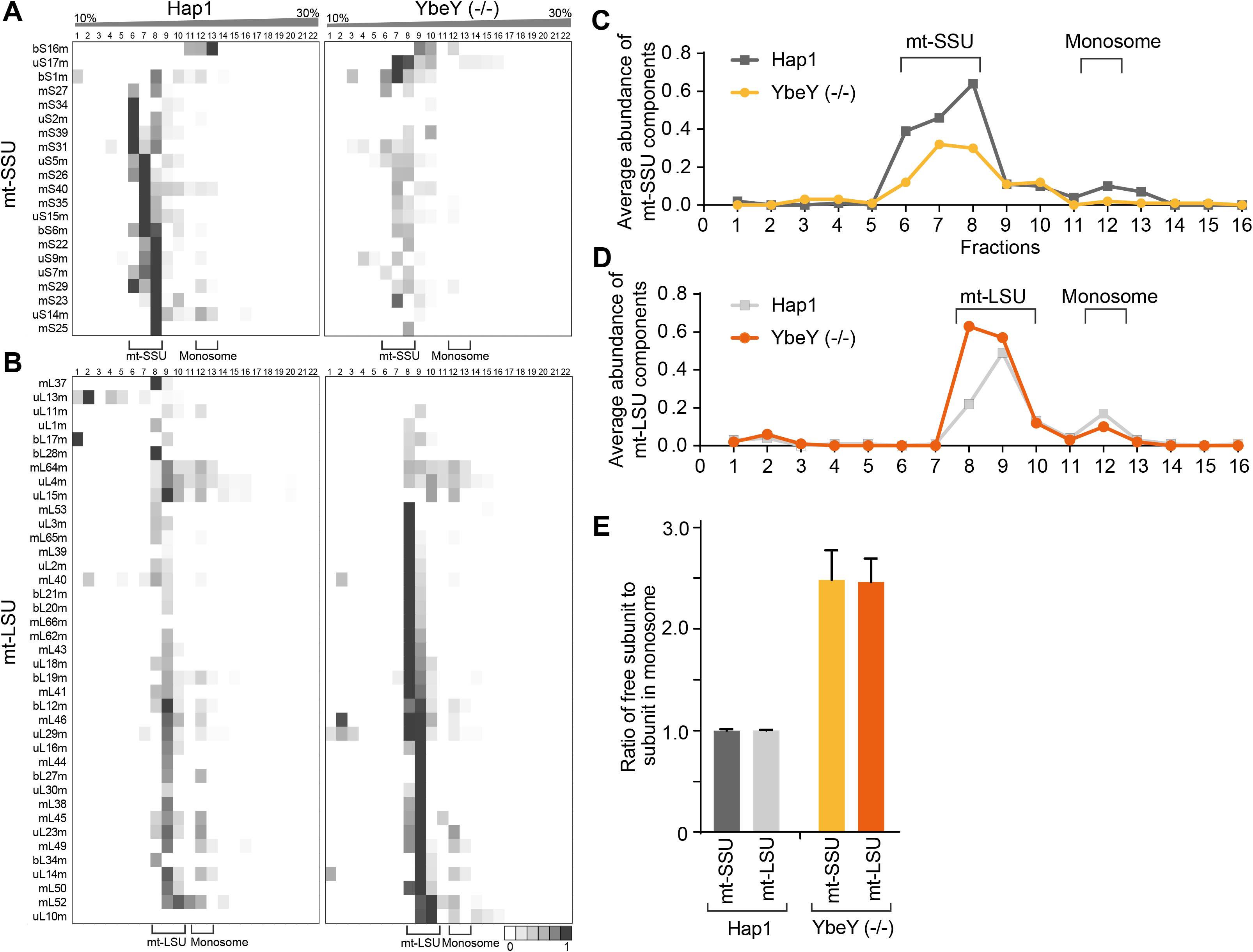
Assembly of mt- SSU is perturbed in YbeY knockout cells. (A, B) Quantitative sucrose gradient fractionation of mtSSU (A) and mtLSU (B). Sucrose gradient fractionation was performed with mitochondria from SILAC labelled parental and YbeY knockout Hap1 cell. The fractions were analysed using mass spectrometry. A heatmap was generated from the intensity of detection of peptides common to both samples. The intensity profile was scaled relative to the maximum intensity. Each row represents a different large subunit component. (C, D) The average abundance of the mt-SSU (C) and mt-LSU (D) components in the sucrose gradient. (E) The proportion of protein components in assembled mt-SSU or mtLSU relative to those found in monosomes was calculated for the Hap1 control cells and the YbeY knockout cells. The abundance of the MRPs in the free mt-SSU (fractions 6-8), the mt-LSU (fractions 8-10) and the monosome (fraction 12) was calculated relative to the total mitoribosomal protein. Then, the proportion of free small or large subunit to the assembled monosome was calculated for the Hap1 parental and YbeY (−/−) cells.

## DISCUSSION

The translation of the mitochondrially-encoded polypeptides requires the coordinated action of numerous factors. Many of these factors act at the RNA level such as cleaving the primary transcript, chemically modifying mt-tRNAs and mt-rRNAs, or chaperoning the mt-rRNA into the correct conformation (52). Here, we have identified YbeY, a mitochondrially-localised protein with a conserved endonucleolytic domain. Deletion of the gene led to decreased steady state levels of the OxPhos complexes which have mitochondrially-encoded components. Moreover, analysis of *de novo* mitochondrial protein synthesis showed that YbeY depletion had a detrimental effect on ribosome activity.

The main catalytic residues involved in endonucleolytic cleavage, are conserved in mammalian YbeY suggesting the conservation of its endonucleolytic function (35). Moreover, YbeY has been shown to be an active RNase in various bacterial species (26,28,37) and in plant chloroplasts (38). In *E. coli*, similar to the mitochondria, the ribosomal RNAs are initially transcribed within a primary transcript and are later released through multiple hierarchical cleavage events. Bacterial YbeY is responsible for the 5’ and 3’ end maturation of the 16S (small subunit rRNA) precursor, the 5’ end maturation of the 23S (large subunit rRNA) precursor and the final steps of 5S rRNA maturation (32). In the mammalian mitochondria, however, the excision of the flanking mt-tRNAs by RNase P and ELAC2 produces the mt-rRNAs ready for mitochondrial assembly. Thus, to investigate if the translation defect was due to errors in RNA processing, we investigated the effect of YbeY depletion on RNA steady state levels. We observed no difference in mRNA and rRNA levels or processing, however, there was a noticeable decrease in the mt-tRNA^Ser(AGY)^ and in mt-tRNA^Leu(UUR)^ to a lesser extent. Analysis of the 5’ and 3’ termini of the tRNAs showed that a severe deficiency in accurate post-transcriptional processing of mt-tRNA^Ser(AGY)^ from the primary transcript in YbeY-deficient cells.

In addition to RNA processing, the bacterial YbeY degrades rRNA from late state ribosomes with defective small subunits and as such is responsible for quality control (26,32). We show that, in the mammalian mitochondria, YbeY is crucial for the integrity of the small subunit. Similar to the bacterial homolog, YbeY may perform its ribosome-associated function through its interaction with the uS11m and ERAL1. Analysis of interactors of the human YbeY showed that it interacts with uS11m and ERAL1, both key components of mt-SSU biogenesis, suggesting a role for YbeY in mitoribosomal biogenesis similar to its role in bacteria. The GTPase ERAL1 is a 12S mt-rRNA chaperone and mt-SSU assembly factor (14). In bacteria, the homolog of YbeY interacts with ribosomal protein S11 (homolog of mitoribosomal component uS11m) and GTPase Era (homolog of 12S mt-rRNA methyltransferase, ERAL1). However, the interaction with Era is facilitated indirectly through its interaction with S11, which explains the weak interaction of ERAL1 with YbeY. A disruption of this YbeY-S11 interaction in bacteria led to an accumulation of 16S small subunit rRNA precursors and a resultant increase in susceptibility to stress (51). The residues involved with the YbeY-S11 interaction are conserved in human YbeY (51). Therefore, this suggests that, similar to its bacterial homolog, YbeY associates with ERAL1 indirectly through uS11m. Both ERAL1 and its bacterial homolog Era binds stem-loop structures at the 3’ end of the SSU rRNA (14,15). In mammalian 12S mt-rRNA, this stem-loop region contains two adenines that are methylated by TFB1M, and this modification is necessary for ERAL1 activity (15). The progression of mt-SSU assembly requires the removal and targeted degradation of ERAL1 from this site (53). Structural analysis of the mitochondrial ribosome suggests that, similar to the bacterial ribosome, the uS11m component is assembled at the 3’ end of the SSU mt-rRNA, at the site of ERAL1 interaction.

Mitochondrial uS11m has extensive connections with the 12S mt-rRNA and binds independently to other protein clusters during the assembly process (23). However, previously published pulse-chase quantitative proteomic analysis of small subunit assembly in bacteria and human cells have demonstrated that while the mitochondrial uS11m subunit is an early binding protein, S11 is one of the last proteins to be assembled into the bacterial SSU (23,54). In addition to uS11m, YbeY interactors also included mS26, mL64, uL2 and early-assembly mt-LSU clusters, bL20m-bL21m-mL42-mL43-mL44 and uL4m-mL50 (23). Mapping these components onto the mitoribosomal structures show that these subcomplexes form a continuous belt of proteins around the large subunit and have extensive contacts with the 16S mt-rRNA (**Supplementary Figure S6**). However, these interactions with the mt-LSU subcomplexes are weak and might not be direct interactions, but secondary interactions via rRNA or other proteins.

The assembly of mt-SSU and mt-LSU can be inextricably linked. It has been previously shown that defects in the assembly of the large subunit have negative consequences for the assembly of the small subunit (7,8). The knockout of YbeY leads to a severe depletion of mt-SSU. We speculate that this impaired assembly of mt-SSU is associated with compensatory accumulation of mt-LSU, owing to perturbed monosome formation. This partly explains the loss in mitochondrial translation in YbeY deficient cells. How this compensatory upregulation of mt-LSU is executed and whether or not it involves sensing of the mt-SSU assembly status, need further investigation.

Alongside its role in mitoribosomal biogenesis, we also demonstrate that YbeY is required for the accurate post-transcriptional processing of mt-tRNA^Ser(AGY)^. The residues required for endonucleolytic cleavage are conserved in YbeY. However, there is no indication of endonucleolytic processing of mt-rRNA during mitoribosomal biogenesis. Normally, the 5’ end of RNAs in the mitochondrial primary transcripts is processed by RNase P before the 3’ end processing by RNase Z (3,21). However, previous work has suggested that mt-tRNA^Ser(AGY)^ is not processed completely at the 5’ end by RNase P (55). The authors suggested that the release of the 5’ end of the mt-tRNA is a consequence of the 3’ endonucleolytic processing of the upstream mt-tRNA^His^, while the presence of the downstream mt-tRNA^Leu(UUR)^ was dispensable for mt-tRNA^Ser(AGY)^ 3’ end processing (55). Furthermore, transcriptomic analysis of RNase P-deficient MRPP3 knockout mice show that mt-tRNA^Ser(AGY)^ is not affected by the lack of RNase P (21) while the presence of ELAC2 is required for the processing of mt-tRNA^Ser(AGY)^ as loss of the enzyme leads to accumulation of unprocessed precursors (56). Here, we demonstrate that in the absence of YbeY, steady state levels of mt-tRNA^Ser(AGY)^ are reduced, and the 5’ and to a lesser extend the 3’ ends of mt-tRNA^Ser(AGY)^ are erroneously processed. However, a small proportion of correctly processed mt-tRNAs are still produced and can be used in translation. We suggest that YbeY works in conjunction with, but independently from, ELAC2 to accurately cleave the mt-tRNA at the 3’ end of the upstream mt-tRNA^His^ and at the 3’ end of mt-tRNA^Ser(AGY)^. The mechanism by which this is performed is currently unclear and remains to be elucidated.

Fractionation experiments are consistent with early mt-LSU assembly and a majority of mt-SSU assembly occurring in the nucleoid (mtDNA containing fractions) after which mitoribosome assembly continues in MRGs, the latter being based on co-localisation experiments (12,20,22–24). Our data suggests that YbeY contributes to late assembly of mt-SSU. However, previously published proteomic analysis of nucleoid-associated proteins did not identify YbeY as a constituent of nucleoids (20) and our results indicate that YbeY does not localise to MRGs. The distribution of YbeY in the mitochondrial matrix may be as a result of its multiple functions in mitochondria. Many mitochondrial RNA binding proteins, such as MRPP1, MRPP2 and NSUN4, have multiple independent functions in RNA modification (57–60). Our results have demonstrated that YbeY, similarly performs multiple functions in the mammalian mitochondria. As such, it may act as a link between the post-transcriptional processing of the primary transcript, and mitoribosomal biogenesis and translation. Moreover, our results also highlight a possible connection between the assembly pathways of the large and small mitoribosomal subunits, where the negative effects on the assembly of the small subunit may have a knock-on effect on the large subunit.

## MATERIAL AND METHODS

### Plasmids

The open reading frame of YbeY was purchased as an IMAGE cDNA clone (IRAUp969G02109D; GeneScript Item number: SC1010) from Source Bioscience and cloned into pCDNA5/FRT/TO constructs.

YBEY + flag Primers
Overhang+BamH1+Forward Primer:
ctttcttggatccaatgagtttggtgattagaaatctgcagcg
Overhang+Not1+Flag+Reverse Primer:
ctctccgcggccgcctacttatcgtcgtcatccttgtaatcgctccctccgaagaggcc

### Genome-modified cell lines

For the generation of YbeY deficient HEK293T cell lines, the cells were transiently transfected with CompoZr ZFNs (Sigma-Aldrich) using Cell Line Nucleofector (Lonza), buffer kit V (Lonza) and program A-023. 72 hours after transfection, the cells were single-cloned into 96 well plates and screened using Sanger sequencing to identify indels at the ZFN target site and western blotting. Wild-type Hap1 and YbeY knockout Hap1 cell lines were purchased from Horizon Discovery (Product ID: HZGHC002765c022).

### Cell culture

Cell lines were maintained at 5% CO_2_ and 37 °C in humidified incubators. The HEK293T and HeLa cells were cultured in Dulbecco’s Modified Eagle Medium (DMEM), containing 4.5 g/L glucose, 110 mg/L sodium pyruvate, supplemented with 10% (v/v) foetal bovine serum, 100 U/ml penicillin and 100 μg/ml streptomycin. Hap1 cells were cultured in Iscove’s Modified Dulbecco’s medium (IMDM) containing 4.5 g/L glucose, 110 mg/L sodium pyruvate, supplemented with 10% (v/v) foetal bovine serum, 100 U/ml penicillin and 100 μg/ml streptomycin. IMDM for YbeY knockout Hap1 cells were supplemented with 20% FBS. Flp-In T-Rex HEK293T cells were cultured in supplemented DMEM with additional 15 μg/ml blasticidin and 100 μg/ml Zeocin. After transfection with pcDNA/FRT/TO plasmid, the cells were cultured in supplemented DMEM containing 4.5 g/L glucose, 110 mg/L sodium pyruvate, supplemented with 10% (v/v) tetracycline-free foetal bovine serum, 100 U/ml penicillin and 100 μg/ml streptomycin, 15 μg/ml blasticidin and 50 μg/ml hygromycin.

### Transfection

2 μg of DNA and 3 μl of Lipofectamine 2000 were incubated in 100 μl of optiMEM separately for 5 minutes at room temperature. They were mixed and incubated for a further 10 minutes. Then, they were added to cells in supplemented DMEM and mixed by swirling.

### Immunodetection of proteins

A confluent 9 cm dish of HEK293T or Hap1 cells (or isolated mitochondria) were washed with PBS and pelleted at 1200 rpm for 3 minutes. The pellet was lysed with Lysis buffer (50 mM Tris HCl, 150 mM NaCl, 1 mM EDTA, 1% Triton, 1 × Roche inhibitor tablet) on ice for 15 minutes and centrifuged at 8000 rpm for 5 minutes. The protein concentration of the supernatant was quantified. 25 μg of lysate was boiled at 95 °C with 33.3 % NuPAGE LDS 4x sample buffer containing 200 mM DTT. The boiled solution was run on a 4-12% Bis-Tris NuPage polyacrylamide gel at 200 V for 30 minutes. The proteins were transferred onto a nitrocellulose membrane with the iBlot 2 Dry Blotting System using Protocol ‘P0’. Gels were stained with SimplyBlue Safestain. The membrane was blocked with 5% non-fat milk in PBS-T at room temperature. The membrane was then incubated overnight at 4 °C with primary antibody diluted in 5% milk in PBS-T. The membrane was washed 3 times for 10 minutes with PBS-T. The membrane was incubated with secondary antibody diluted in PBS-T and then washed again as before. Finally, the membrane was developed using ECL and X-ray film.

### Primary antibodies

TOM22 Mouse 1:2000 (Abcam ab10436)
GAPDH Mouse 1:2000 (Abcam ab10436)
YbeY Rabbit 1:500 (Atlas HPA018162)
Flag Mouse 1:1000 (Sigma F3165)
TOM20 1:500 (Santa Cruz sc-11415)
OXPHOS cocktail Mouse 1:1000 (Abcam ab110411)
uL3m Rabbit 1:2000 (MRPL3, Atlas HP043665)
mS40 Rabbit 1:2000 (MRPS18b, Proteintech 16139-1-AP)
uS17m Rabbit 1:2000 (MRPS17, Proteintech 18881-1-AP)
mS35 Rabbit 1:1000 (MRPS35, Proteintech 16457-1-AP)
bL12m Rabbit 1:2000 (MRPL12, Proteintech 14795-1-AP)
NDUFS1 Rabbit 1:2500 (Abcam ab169540)

### Secondary antibodies

Goat anti-rabbit IgG HRP 1:1000 (Promega W4011)
Goat anti-mouse IgG HRP 1:2000 (Promega W4021)

### Mitochondrial Isolation and cell fractionation

Mitochondria were isolated using a cell homogeniser (isobiotec) as described by (61). For cell fractionation, the mitochondrial isolation protocol using a Dounce homogeniser adapted from (62) was used. Briefly, 15 confluent 15 cm dishes of HEK293T cells were washed with 1 × PBS and pelleted. The cell pellet was resuspended in 15 ml of mitobuffer (0.6 M mannitol, 10 mM Tris HCl pH 7.4, 1 mM EDTA) containing 0.1% BSA. 100 μl of the cell suspension was kept aside as the ‘total fraction’. The suspension was homogenised in a Dounce homogeniser (15 strokes). The homogenised suspension was centrifuged at 400 g for 10 minutes. The pellet was resuspended in mitobuffer containing BSA and re-homogenised. The supernatant was used as the ‘debris fraction’. The supernatant was centrifuged at 400 g for 5 minutes. The supernatant from this step was centrifuged at 11,000 g. The supernatant from this step was kept aside as the ‘cytoplasmic fraction’. The pellet was resuspended in 1 ml of mitobuffer containing 50 U/ml Benzonase, incubated on ice for 15 minutes and centrifuged at 11,000 g for 10 minutes. The mitochondrial pellet was resuspended in 120 μl of mitobuffer and divided into two aliquots. The first aliquot was the mitochondrial fraction. The second aliquot was incubated with 4 μg/mg of Proteinase K for 30 minutes on ice (proteinase K fraction). The aliquots were centrifuged at 11,000 g for 5 minutes. The pellet was lysed, and all fractions were analysed by western blotting.

### Immunocytochemistry

The cells were grown on coverslips in a well of a 6-well plate. 24 hours after transfection, the cells were washed with PBS and then incubated with 4 % formaldehyde diluted in PBS for 15 minutes. The cells were then washed with PBS and then permeabilised for 5 minutes with 0.1% Triton-X diluted in PBS. The cells were incubated for 1 hour in PBSS (5% FBS v/v, 95% PBS) and then for 1.5 hours in primary antibody diluted in PBSS. The cells were washed three times for 5 minutes in PBSS. Secondary antibody, diluted in PBSS, was added to the cells and incubated for 1 hour. Following the incubation, the cells were washed (5 minutes each) twice in PBSS and once in PBS and mounted in Vectashield mounting medium. The cells were visualised by fluorescence microscopy using a Zeiss LSM 510 META confocal microscope.

### Primary antibodies

Flag 1:200 (Sigma F3165)
Tom20 1:500 (Santa Cruz sc-11415)

### Secondary antibodies

Alexa Fluor 488-conjugated goat anti-mouse 1:1000 (Molecular Probes A1101)
Alexa Fluor 594-conjugated goat anti-rabbit 1:1000 (Molecular Probes A11037)

### Circularisation RT-PCR sequencing

For the analysis of the 5’ and 3’ ends of certain mt-tRNAs and mt-rRNAs, the protocol was adapted from (48). In brief, the RNA was circularised and reverse transcribed across the ligated junction. PCR was used to amplify the product, which was then cloned using a Zero Blunt TOPO PCR cloning kit and sequenced using an M13 forward universal sequencing primer.

### ^35^S-methionine metabolic labelling of *de novo* protein synthesis

Newly synthesised mitochondrial proteins in Hap1 and HEK293T cell lines were labelled with ^35^S-labelled methionine and analysed by autoradiography as described by (63).

### Quantitative immunoprecipitation

Doxycycline-inducible YbeY.FLAG-expressing cells were grown in heavy (^15^N and ^13^C labelled Arg and Lys containing medium) and light (^14^N and ^12^C Arg and Lys (Sigma Aldrich) containing medium) SILAC medium. Cells were differentially induced with 100 ng/ml doxycycline for 24 hours. The four conditions (heavy induced, heavy uninduced, light induced, light uninduced) were harvested, pelleted at 1200 rpm for 3 minutes, washed with PBS and re-pelleted. The cells were quantified and mixed in equal proportion (heavy induced with light uninduced, and heavy uninduced with light induced). Mitochondria were isolated (see Mitochondrial isolation section) from the cell suspension. Mitochondria were lysed and immunoaffinity purification was carried out with FLAG M2 affinity gel (Sigma-Aldrich) following manufacturer’s instructions. Elution was performed with FLAG peptide. The interactors were measured using mass spectrometry and the interactors were separated into ‘induced’ and ‘uninduced’ based on isotope labelling (see Mass spectrometry and analysis of SILAC samples section). After protein concentration and labelling were computationally identified. The reciprocal of the protein orientation was calculated for the reverse labelling orientation. Ratios for both the samples and a base two logarithm were calculated. The data was plotted on an X-Y scatter plot.

### Quantitative isokinetic sucrose gradient analysis of mitoribosomes

6 × 15 cm dishes of Hap1 control and YbeY (−/−) Hap1 cells were grown in heavy (^15^N and ^13^C labelled Arg and Lys containing medium) and light (^14^N and ^12^C Arg and Lys (Sigma Aldrich) containing medium) SILAC medium respectively. The cells pelleted at 1200 rpm for 3 minutes, washed with PBS and re-pelleted. The cells were resuspended in 20 ml of PBS, quantified using the Pierce BCA protein assay kit as per manufacturer’s instructions and the two cell suspensions were mixed in equal proportions. The cells were pelleted and mitochondria were isolated (see Mitochondrial isolation section). The mitochondrial pellet was incubated on ice with 100 μl of lysis buffer for 15 minutes and centrifuged at 8000 rpm for 5 minutes at 4 °C. 750 μg of the lysate was run on a 10-30% isokinetic sucrose gradient as described in (63). 100 μl fractions were taken and run 4 mm into a 4-12% Bis-Tris polyacrylamide gel to remove the sucrose. The gel was stained for 1 hour with SimplyBlue Safestain and destained for 1 hour (20% methanol, 10% acetic acid). The stained 4 mm region was cut and analysed by mass spectrometry. The Hap1 and YbeY(−/−) proteins were separated by their SILAC labelling.

### Mass spectrometry and analysis of SILAC samples

The peptides were analysed with LC-MS/MS on a Thermo Scientific Oribitrap LTQ XL spectrometer with a nanospray inlet interface coupled with Proxeon EasynLC nano chromatography. The internal ‘lock mass’ standard of 445.120 m/z was used for accurate mass measurements. Samples were run from the earliest fraction to the latest without intervening washes to maintain chromatographic conditions between washes. To identify peptides, resulting data files were processed using Proteome Discoverer software (ThermoFisher Scientific). The resulting spectra from the RAW file were submitted to the in-house Mascot server and peptides were assigned using the Uniprot *Homo sapiens* protein database, specifying parameters for use of trypsin digest of the gel pieces, allowing for the possibility of one uncleaved lysine and arginine residue per peptide, allowing for peptides to carry a deviation of 10 ppm around calculated peptide values to allow matches with experimentally observed values, and fragment size was set at 0.5 Da. A multiconsensus report was generated following the quantification of each identified peptide using Protein Discoverer, allowing the relative quantification of proteins to be obtained from the relative quantification of peptides. The mean value of the three most abundant peptides for each protein was used as a representative value for that protein. This was performed for fraction and a profile for each protein for each fraction was determined.

For SILAC samples, proteins were identified using the ANDROMEDA algorithm of the MAXQUANT software package by comparing the proteins to the human UniProt database. The peptide information from both sets were combined and filtered using the PERSEUS software, removing proteins only identified by site, that matched a decoy peptide database and contaminating peptides. Proteins identified by two or fewer peptides were removed. Proteins identified from only one of the two cell lines were removed from the final protein dataset. Heavy label incorporation was performed by manually processing peptide information of the heavy-only sample by MAXQUANT. The maximum intensity for each of the proteins in the two samples was marked as the maximum value and the remaining intensity profile of that protein was scales proportional to this value.

### RNA isolation and northern blotting

RNA was extracted from HEK293T and Hap1 cell lines with TRIzol reagent (ThermoFisher Scientific) according to manufacturer’s instructions. Northern blot was performed as described by (62) with minor modifications. Total RNA was resolved on a 6% or a 15% urea-PAGE gel (for mt-tRNAs). After wet transfer of the RNA to a nylon membrane (Magnoprobe, 0.45 μ; GE Healthcare or Genescreenplus, NEN DuPont) in 2 × SSC (150 nM NaCl, 15 mM sodium citrate pH 7.0), the membrane was subjected to UV-crosslinking (0.12 J). High resolution tRNA gels were dry blotted using the ‘rapid transfer method’ (Reference). The membranes were hybridised overnight at 65 °C with radioactive probes in 7% SDS and 0.25 M sodium phosphate buffer (pH 7.6), washed with SSC (containing 0.1% SDS) five times for 20 minutes. The membrane was exposed to a storage phosphor screen (GE Healthcare), visualised using a Typhoon phosphorimaging system and quantified using ImageJ (http://imagej.nih.gov/ij).

## ACKNOWLEDGEMENTS

We are grateful to Michael Harbour, Shujing Ding and Ian M. Fearnley for their help in the proteomic analysis.

## Supplementary Information

**Supplementary Figure S1 (related to Figure 1).**
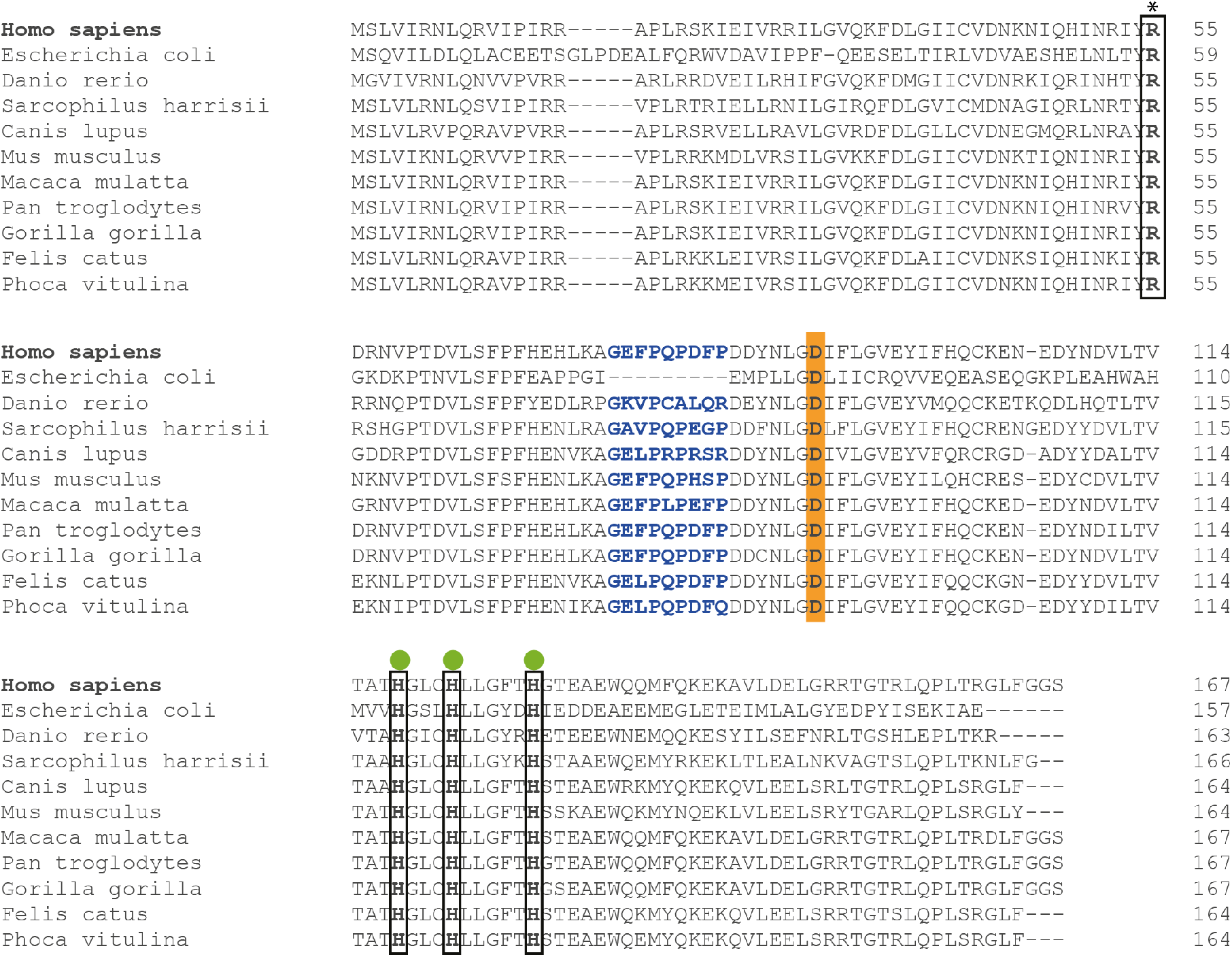
Sequence conservation in YbeY. The YbeY amino acid sequences of various species were aligned using Clustal Omega tool. The catalytically-important RNA-binding arginine is indicated with a box and an asterisk above. Green circles indicate histidine residues from the zinc ion-binding H3H5XH motif. Asp90 (equivalent of Asp85 in *E. coli*), found in the beta-sheet outside the active site and required for the interaction of YbeY with the ribosomal SSU component S11 is indicated in orange. Eukaryote-specific insertion is indicated in blue.

**Supplementary Figure S2 (related to Figure 1).**
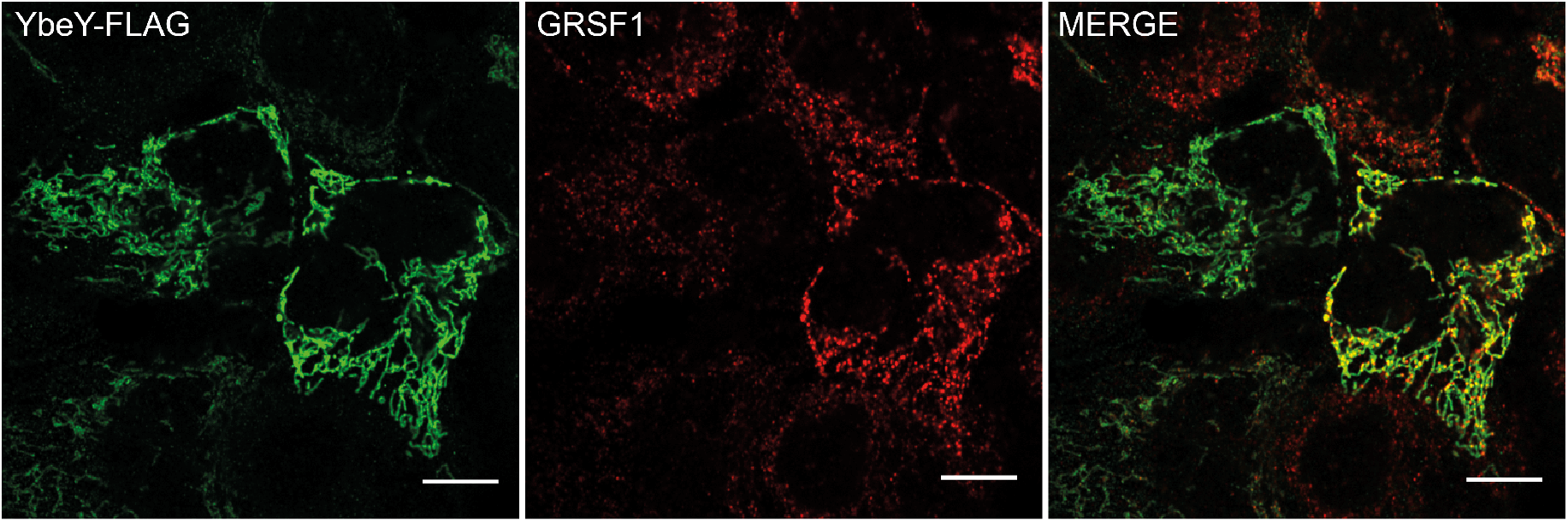
Investigating YbeY colocalization in mitochondrial RNA granules using immunocytochemistry. Flag-tagged YbeY was transiently expressed in HeLa cells and visualised using anti-flag antibodies and visualised using secondary antibodies conjugated with Alexa fluor 488 (green, left). RNA granules were stained using antibodies against GRSF1 (red, middle). Co-localisation is shown in yellow. The white scale bar at the bottom depicts 10 μm.

**Supplementary figure S3 (related to Figure 2).**
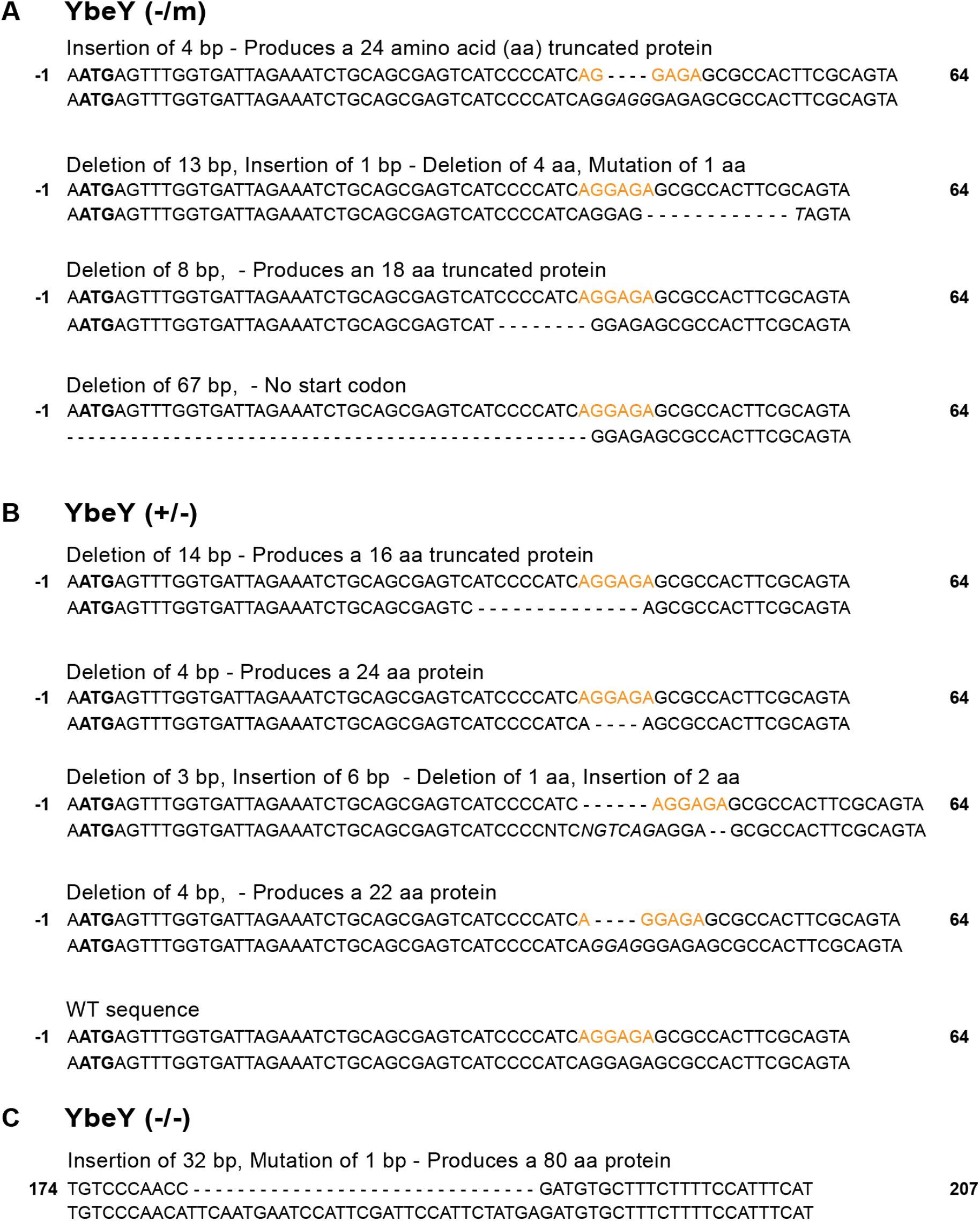
Genome modifications of the *YbeY* locus. PCR products from candidates were cloned into pCR4 vectors using Zero Blunt TOPO cloning kit and bacterial colonies were picked and sent for Sanger sequencing. Indels at the ZFN target sites (depicted in red) are shown. (A) Four alleles of YbeY were identified in YbeY(−/m) clone (B) Five alleles of YbeY were identified in YbeY(+/−) clone. The wild type sequence is above in the pair of sequences. The Start codon is in bold. The zinc finger nuclease target site is coloured orange. (C) A single allele is present in the YbeY(−/−) Hap1 cells.

**Supplementary Figure 4 (related to Figure 2).**
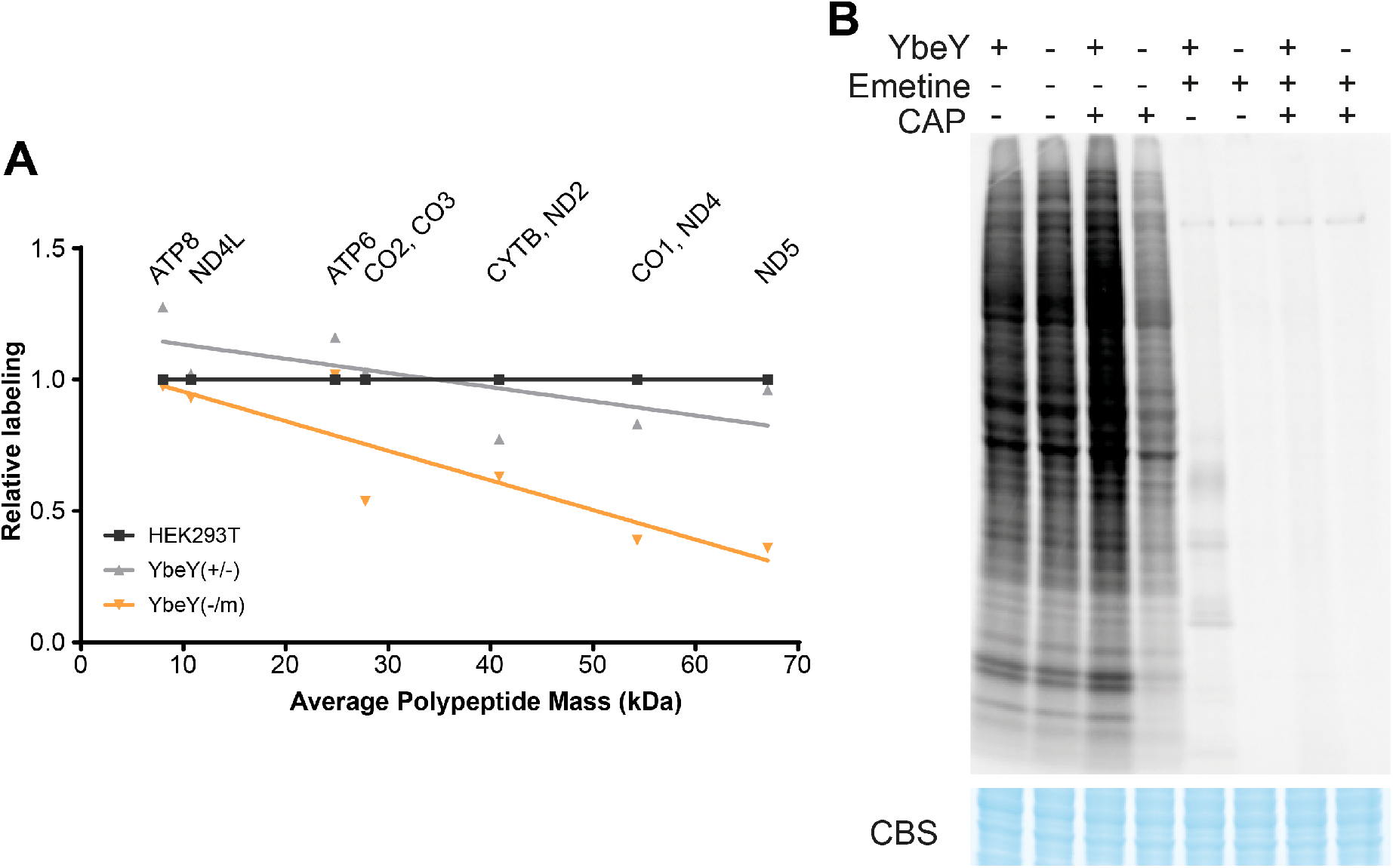
Mitochondrial translation in YbeY-deficient cells. (A) Quantification of mitochondrial translation products relative to the expected size of the polypeptide. Relative metabolic labelling of the polypeptides was quantified from the intensity of the bands using ImageJ, measured relative to the control wild type sample. The bands specific for CO2 and CO3, CYTB and ND2, CO1 and ND4 were quantified together due to their proximity on the gel. The average mass of both proteins was calculated. The line represents the linear regression. YbeY(+/−) R^2^=0.46, YbeY(−/m) R^2^=0.77 (B) Metabolic labelling of mitochondrial translation products in Hap1 parental cells and YbeY knockout cells in the presence and absence of cytoplasmic inhibitor, emetine, and mitochondrial inhibitor, chloramphenicol. Coomassie Blue staining (CBS) was used as loading control.

**Supplementary Figure 5 (related to Figure 3).**
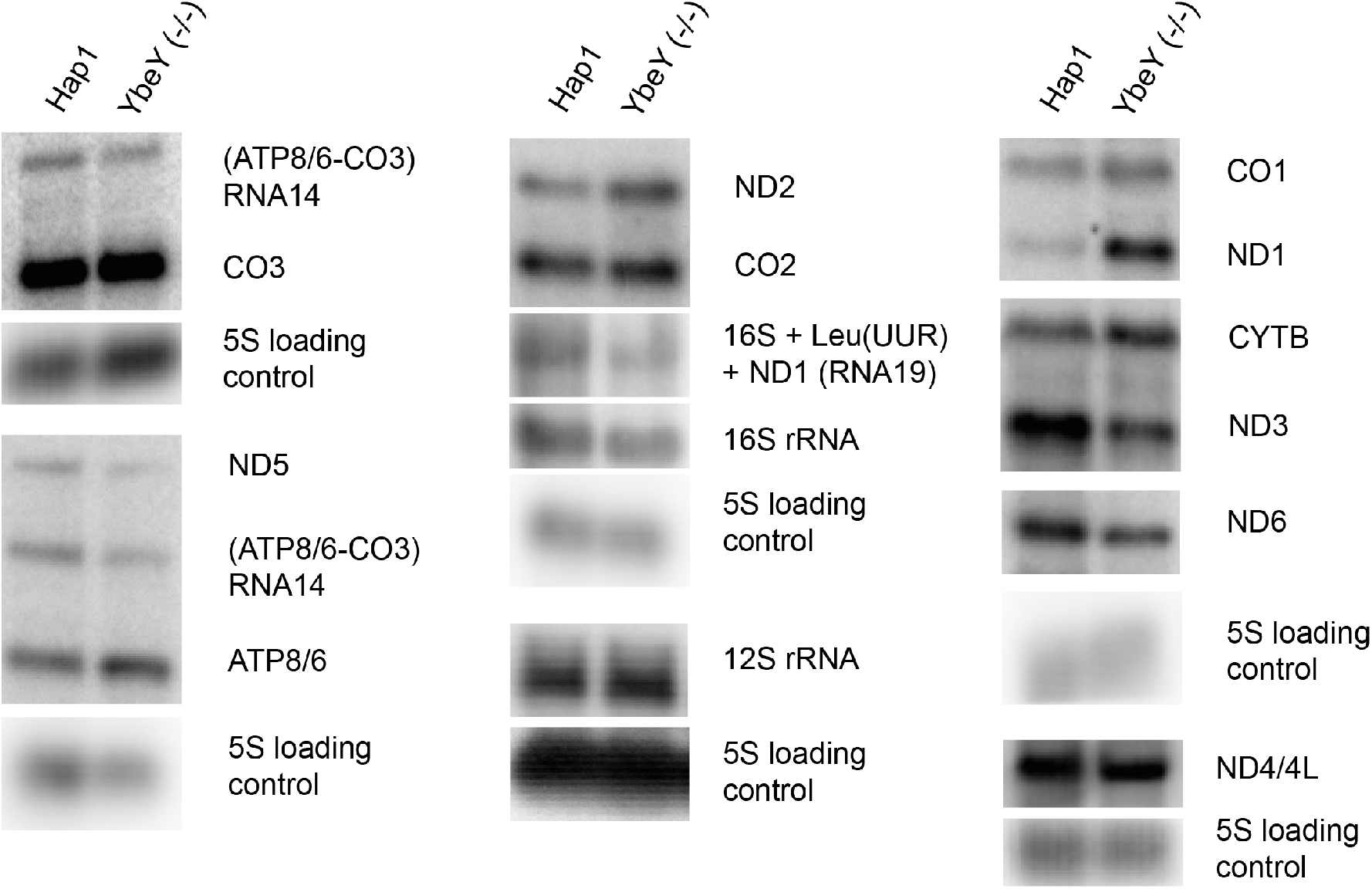
Northern blot analysis of mitochondrial transcripts. Northern blot of mitochondrial mRNAs and rRNAs in YbeY knockout Hap1 cells. 5S RNA was used as a loading control.

**Supplementary Figure S6 (related to Figure 5).**
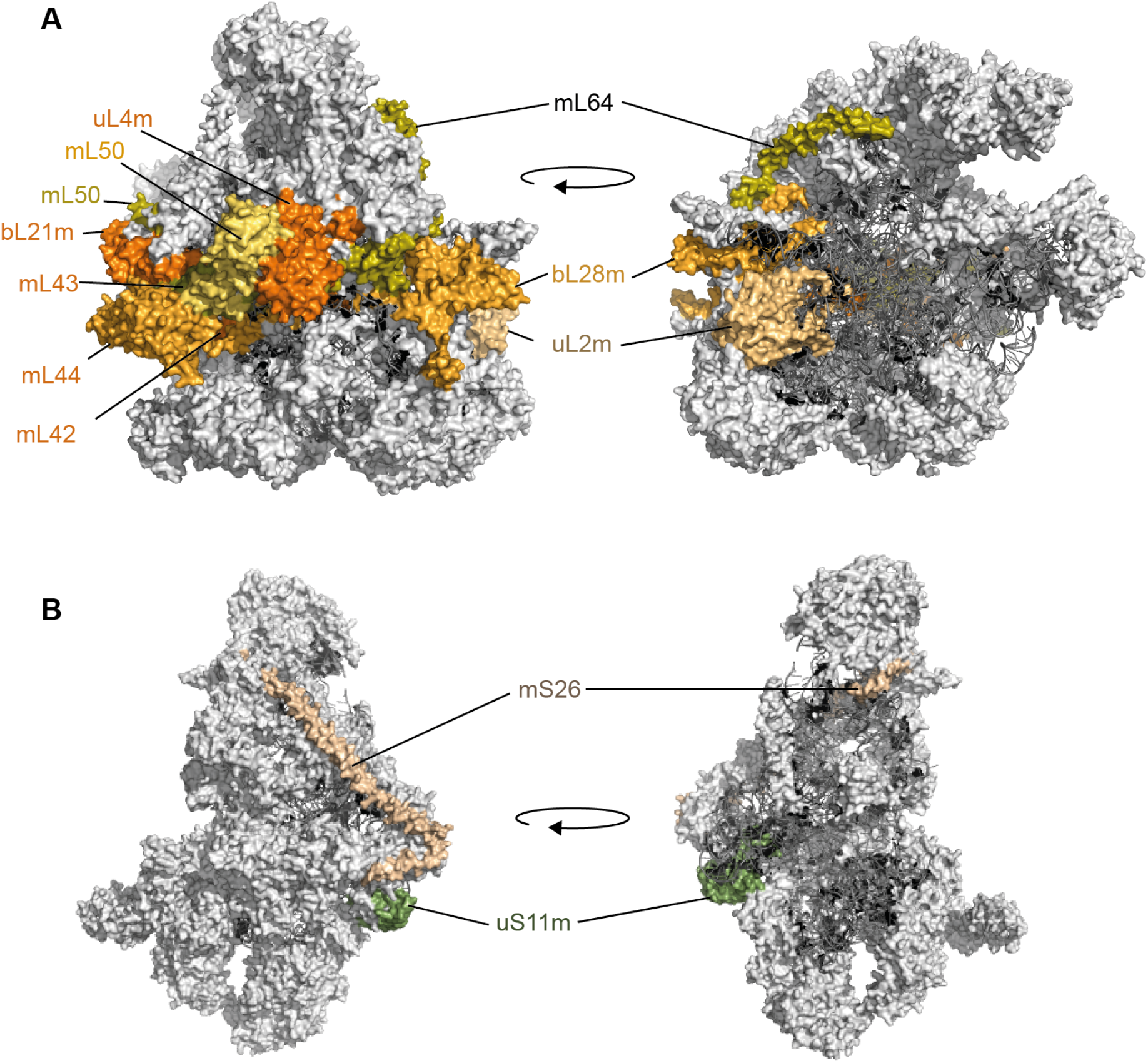
YbeY interactors on the mitochondrial ribosome. Mitoribosomal proteins (light grey), rRNA (dark grey). (A) mt-LSU (B) mt-SSU (PDB: 3J9M) The bL20m-bL21m-mL42-mL43-mL44 subcomplex (orange labels) and uL4m-mL50 subcomplex (green labels).

**Supplementary Figure S7 (related to Figure 6).**
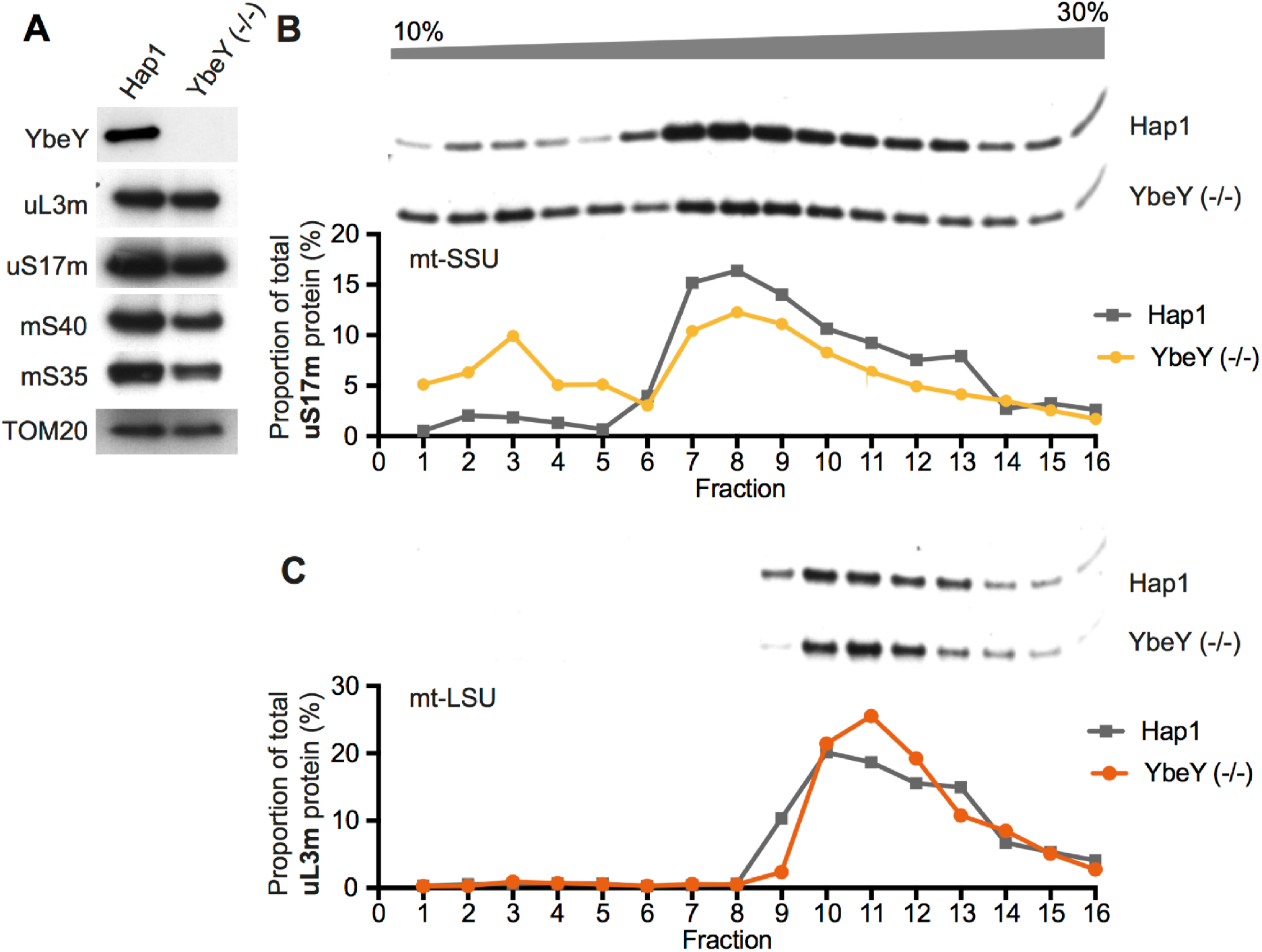
Mitochondrial SSU assembly is reduced in YbeY knockout cells. (A) Analysis of steady state levels of mitoribosomal proteins and endogenous YbeY in Hap1 parental and YbeY knockout cells by western blot. TOM22 was used as a loading control (B, C) Analysis of mitochondrial subassemblies of the large subunit (uL3m) and the small subunit (uS17m) using isokinetic sucrose gradient fractionation and western blotting.

